## Supplementary material for "Learning orientation-invariant representations enables accurate and robust morphologic profiling of cells and organelles": All supplementary notes

### Supplementary 1

This supplementary section supports the second results section and the relevant Methods sections. The topics are: image orientation, pre-alignment, and their impact on embedding accuracy.

#### Workflow of standard pre-alignment algorithms

**Supplementary Fig 1a: Workflow of standard pre-alignment algorithms**
See Methods for procedure details and citations. All images (left box) are rotated so that its 'major axis' aligns with the y-axis (second box). Since the object can be flipped in the x- or y-axes or both, there are four valid flips that still have aligned major axes. To resolve this, each of the four flips is computed (third box), and the 'best flip alignment' is chosen as the final orientation (right box). The criteria for 'best flip alignment' is having highest cross-correlation with a reference image. and the reference image is the mean image in pixel space (bottom box). The flip-alignment stage is repeated until the cross-correlation with the mean image converges (corresponding to a consistent set of flips), which in our experiments took fewer than 30 iterations.

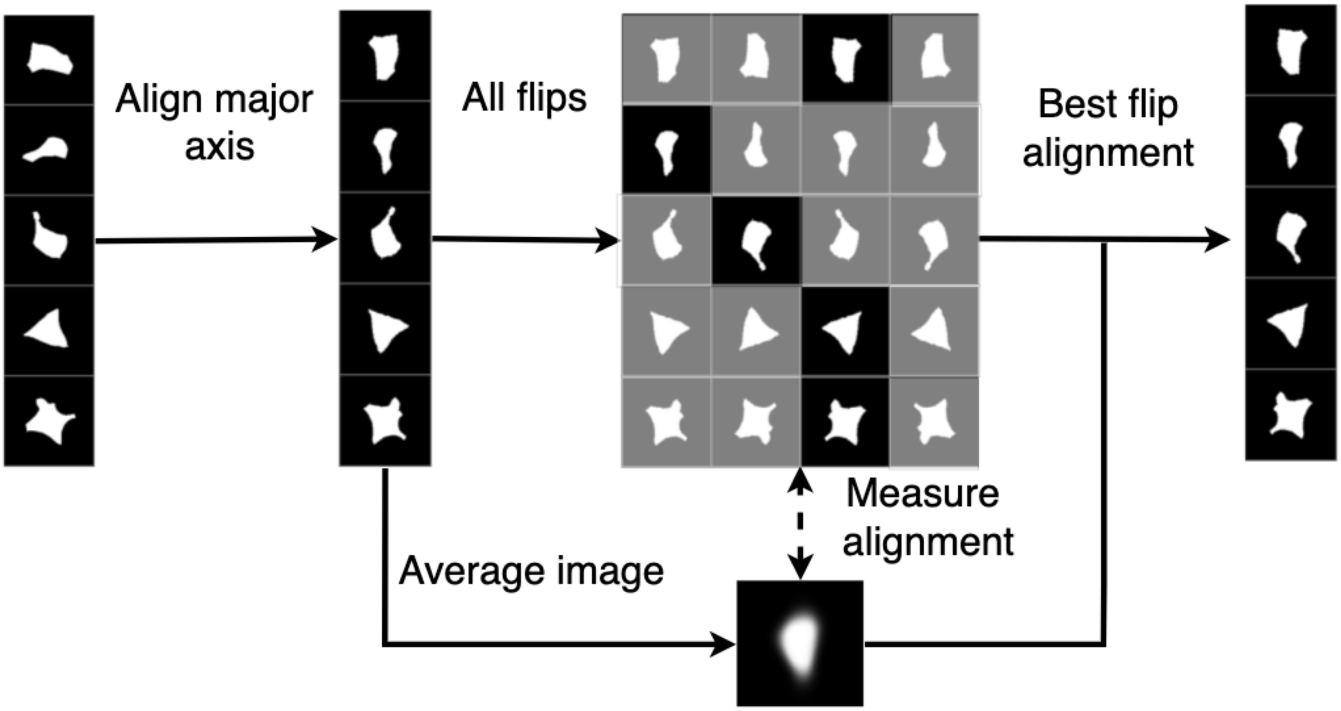


#### Embedding error margin for varying threshold

**Supplementary Supplementary Fig 1b: embedding error margin for varying threshold**
The 'embedding error' score in Fig.2c-d depends on a choice of threshold, which we chose as t=100. Here we plot the embedding error for O2-VAE and prealign-VAE at many choices of `k`

and for each cell line. This shows that prealign-VAE has more embedding errors than O2-VAE for a wide range of choices for t, so the Fig.2c-d results are not an artifact of the choice of t.

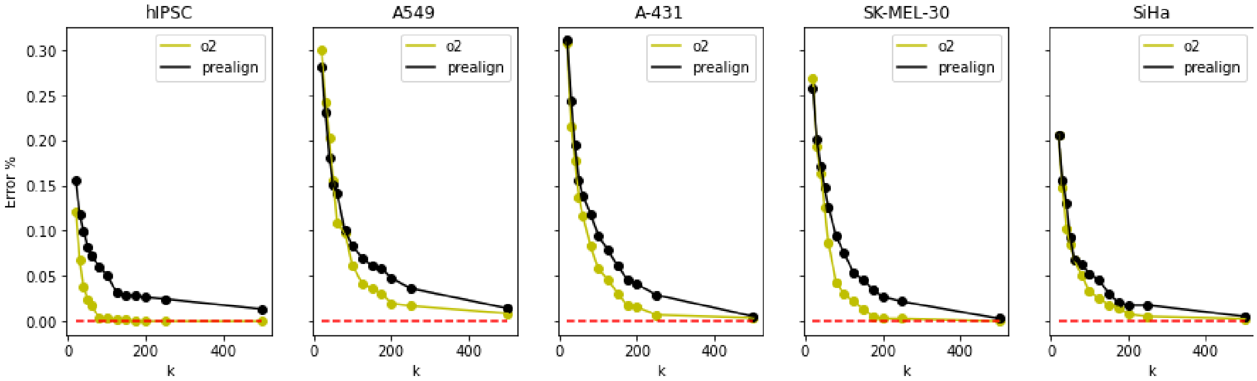


#### Supporting discussion: pre-alignment errors cause embedding errors, and it is difficult to fix them

We first investigate embedding errors for prealign-VAE and establish that they are likely caused by pre-alignment errors.

**Supplementary Figure 1c: pre-alignment errors explain the gap in prealign-VAE vs O2 error rates**
For each test dataset (Allen hiPSC and four human protein atlas cell lines), this displays sample image pairs that are embedding errors for prealign-VAE but not O2-VAE. For each, column 1 and 2 are pairs, and column 3 and 4 are pairs.

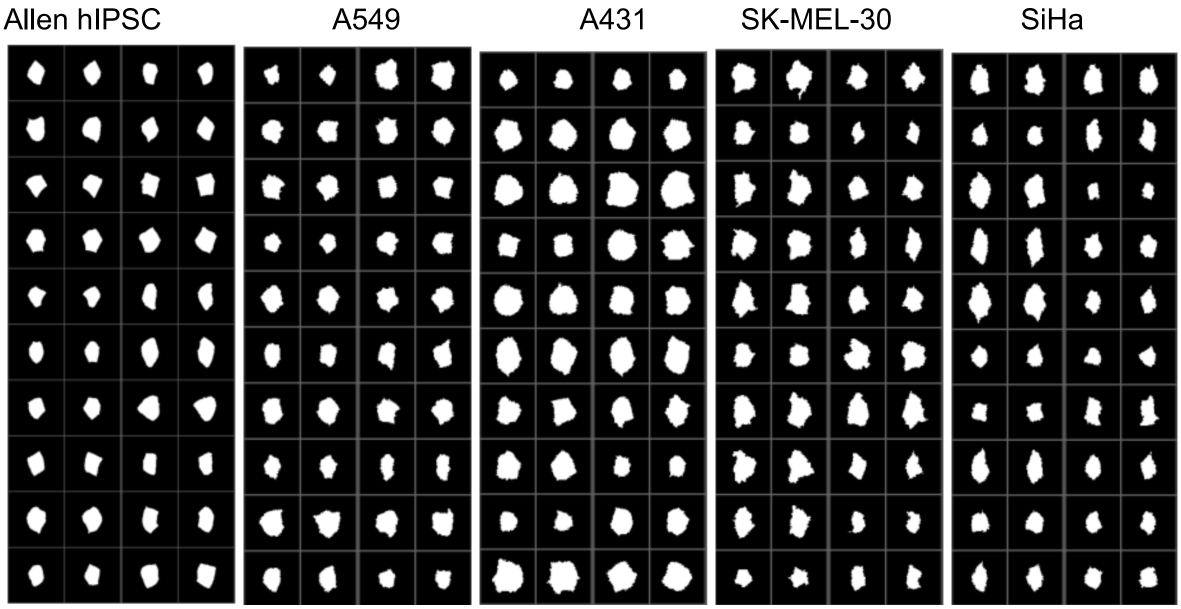


In Results, we claimed that prealign-VAE has higher embedding errors because of image pairs having bad alignments, and that is supported by Supplementary Figure 1c. Most of these pairs are similar in shape, but they are poorly aligned with each other. Only a minority of these pairs are actually well aligned.

**Supplementary Figure 1d: Embedding errors and their issues with pre-alignment**
Investigating 16 example pairs from the hiPSC dataset that are "embedding errors" for the prealign-VAE (which is just a regular VAE where images are pre-aligned, as explained at the start of this Supplementary). Each row is a pair and we label them with an index between 0 and 15 above the first image in the row. The 1st and 2nd columns are the two objects after aligning them optimally, and the 3rd column is the residual plot (1st column minus 2nd column in pixel space). The 4th and 5th columns show the images in their orientation after applying the pre-alignment algorithm, and the 6th column is their residual (4th column minus 5th column in pixel space). The 7th column shows the second object in its optimal alignment with respect to the first object. For the 7th column, we also draw a line corresponding to its major axis and write the angle between this and the y-axis above the image. Since the first object has its major-axis on the y-axis, this also measures the angle between first and second object major axes when they are aligned with each other.

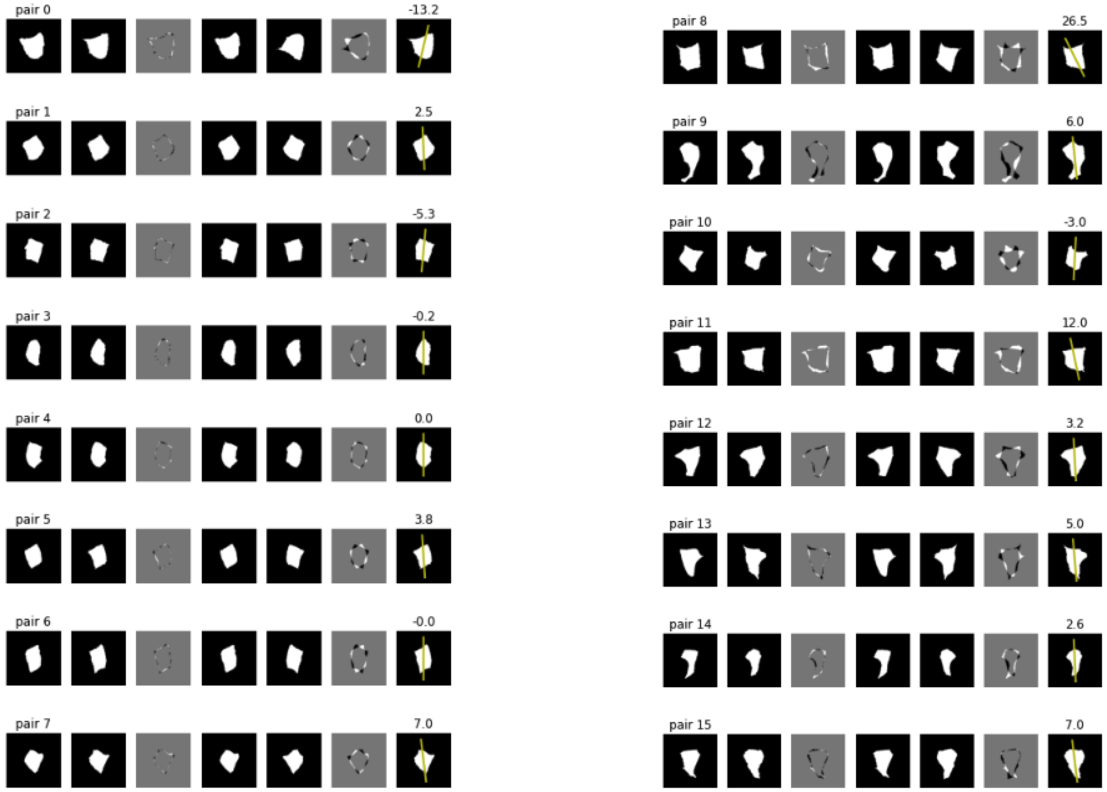


Before explaining insights from this figure, we need one more figure:

**Supplementary Figure 1e: flip alignment decision for images with embedding errors**
Recall that the pre-alignment procedure aligns the major-axis and then flips the image to best align with a reference cell (the reference cell is the image mean in the dataset). Below, each row is one of the images from Supplementary Figure 1c in each of the four possible flip orientations. The image shows the residual compared to the mean cell, and the number above is the

absolute pixel error over the image. The 1st column of each row is the orientation with minimal pixel error - this is the orientation chosen by the pre-alignment algorithm.

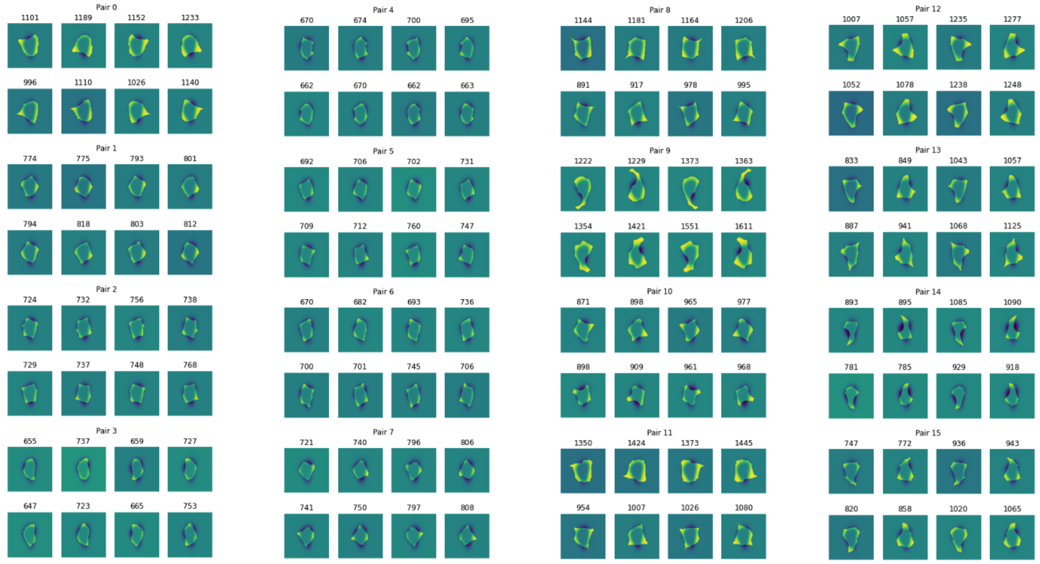


The goal of pre-alignment is to ensure that any image pairs with similar shapes (like all these examples) will be oriented in the same way. For these embedding errors, it's clear that most of the images after pre-alignment are mis-aligned with each other. Supplementary Figure 1c shows this: columns 4 and 5 are the pre-aligned objects, and column 7 shows the orientation of the second image if properly aligned with the first image.

**Failure modes of pre-alignment**

Supplementary Figures 1d and 1e highlight two major failure cases.

Case 1 is *major-axis mis-alignment*. The objects in their optimal alignment have different major-axis angles. In Supplementary Figure 1d, column 4 is the first object after pre-alignment, and column 7 is the second object aligned with the first object with its major axis annotated. This failure case has the major axis that is far from the y-axis (example images 0, 7, 8). The best pairwise alignment is therefore not possible because pre-alignment always forces the major axis to be aligned with the major axis. So this constraint makes pre-alignment computationally tractable, but also introduces this failure case.

Case 2 is *unstable flip alignment*. Image orientation is unstable in the sense that the pre-aligned image and its flipped version have very similar cross-correlation with respect to the mean cell. A slightly perturbed version of the same image may therefore be flipped by the pre-alignment procedure. All examples above that are not case 1 (i.e. examples with a small angle in column 7) are examples of unstable flip. Supplementary Fig 1e shows that flipped versions of the image have very similar error. This is the more common case.

**Commentary**

One natural approach is to add new heuristics to the pre-alignment algorithm to address the 2 failure modes just described.

We experimented with changes to the pre-alignment algorithm to address the failure case 2. We hypothesised that some objects have similar flip cross-correlations with the mean cell because they are very small (or very large) compared to the mean cell. The reasoning was that for very small (or large) cells, the contour details of the cell were fully contained (or fully outside the bounds of) the mean cell, so the correlation error would not change when flopping. The solution was to scale-normalise all objects before doing flip-alignment.

But this modified version of pre-alignment still had failure cases. This suggested that finding heuristic fixes to pre-alignment is very challenging. Heuristic fixes that help with one dataset may not help with another one. We hypothesise that it is difficult to find a pre-alignment algorithm that is simple, while working robustly across many datasets.

Looking into the failure cases specifically, Failure case 2 appears because the dataset is diverse, so there is no single image that can serve as a good reference for all input images. Failure case 1 is very difficult to overcome without a complete change of approach. The choice of aligning the major axis with the y-axis constrains the search space of image orientations. If we don't do this constraint, the optimization would become computationally intractable.

In summary, global pre-alignment is hard because (i) major axis alignment - which is necessary to make the algorithm computationally tractable - causes some similar pairs to be misaligned; and (ii) it is difficult to find a good reference shape to do registration for a diverse dataset. We claim that these issues are very hard to overcome.

Finally we make the point that these errors are present for segmentations of cells, which is a simple setting (the only simpler one would be segmentation of nuclei). In settings, such as joint cell and nucleus segmentations or grayscale images, pre-alignment is probably even more challenging.

#### Methods for finding similar-object-pairs in embedding error experiments

First we justify the similarity metric proposed in the Methods, especially the fact that we raised the normalizer to the power of 1/2. We want a metric that can identify pairs that are very likely to have similar shape. The RMSE is one approximation:

$$RMSE(x,y) = \left( \frac{1}{m \cdot n} \sum_{i,j} (x_{i,j} - x_{i,j})^2 \right)$$

But we observe that if the scale of an object increases, the same (relative) perturbation will cause more pixels to change, increasing this score. In RMSE, the sum of squared differences is scaled by the number of image pixels: $m \cdot n$. To avoid the scale bias, we replace this scalar by

the size of the image. For binary masks, this is the number of nonzero pixels, which we write with the function $s(\cdot)$. We scale by whichever image is largest: $max\{s(x), s(y)\}$.
Replacing the scaling terms $m \cdot n$ with $max\{s(x), s(y)\}$ is mathematically equivalent to dividing the RMSE with by the scalar:

$$C = \left(\frac{m \cdot n}{max\{s(x), s(y)\}}\right)^{\frac{1}{2}}$$

And dividing by this scalar gives the $NRMSE$ introduced in the Methods. (Actually we remove the $\sqrt{m \cdot n}$ term since this is the same for all images, and so it does not effect the relative ordering of $NRMSE$ scores, which is all we need).

Next, we show evidence that “high-confidence-similar-pairs” really are semantically similar with the below figure.

**Supplementary Figure 1f: Sample ‘high-confidence-similar-pairs’ for embedding error experiments**
We show a random sampling of pairs thresholded to be ‘similar with high confidence’ (column 1 and 2 are pairs, column 3 and 5 are pairs, etc). In experiments we try a range of thresholds, so for each dataset below, the left plot is a sample of images when using the smallest threshold, and the right plot is using the largest threshold. They are plotted in the orientation after doing pre-alignment. Cell lines are: **a** hiPSCs, **b** HPA A 549, **c** HPA A 431, **d** SK MEL 30, **e** SiHa.

**(a)**

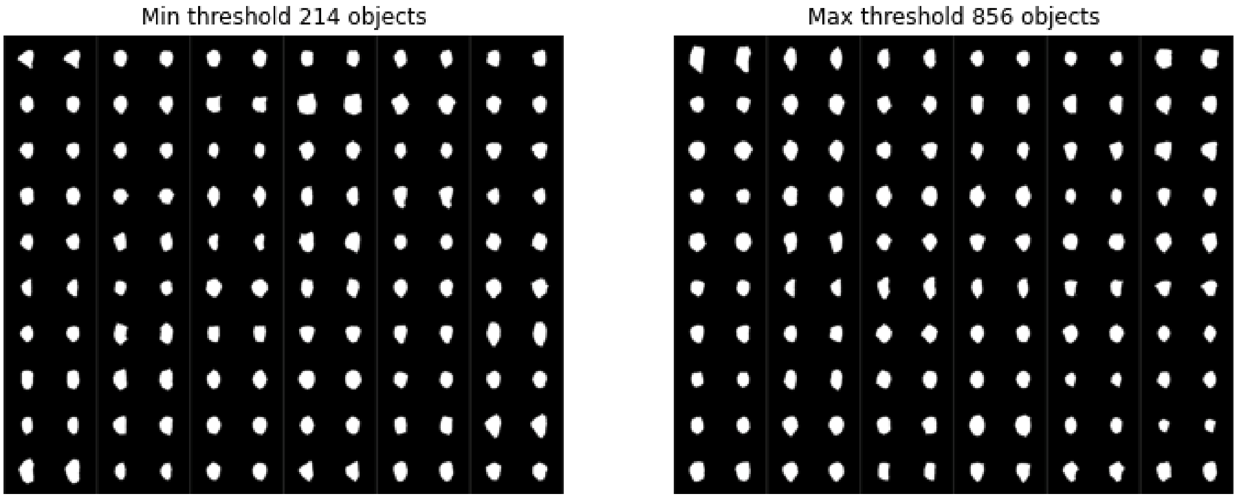

(b)

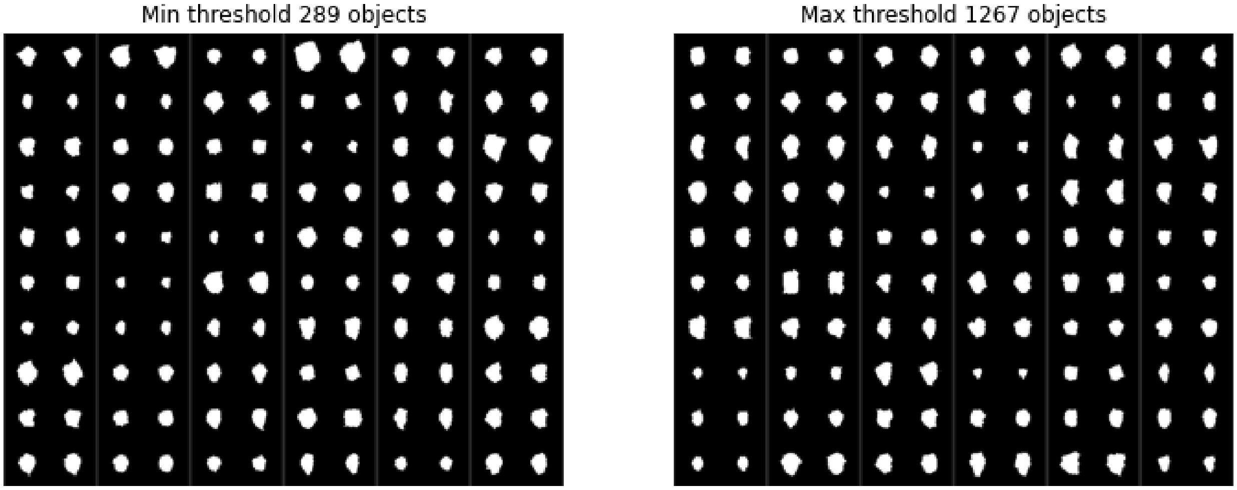

Min threshold 289 objects
Max threshold 1267 objects


(c)

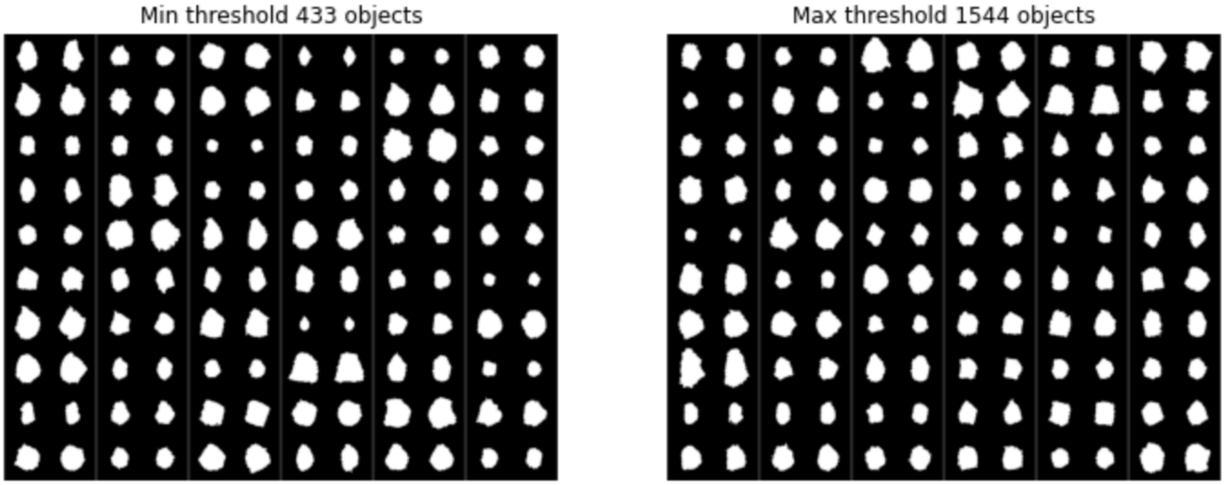

Min threshold 433 objects
Max threshold 1544 objects


(d)

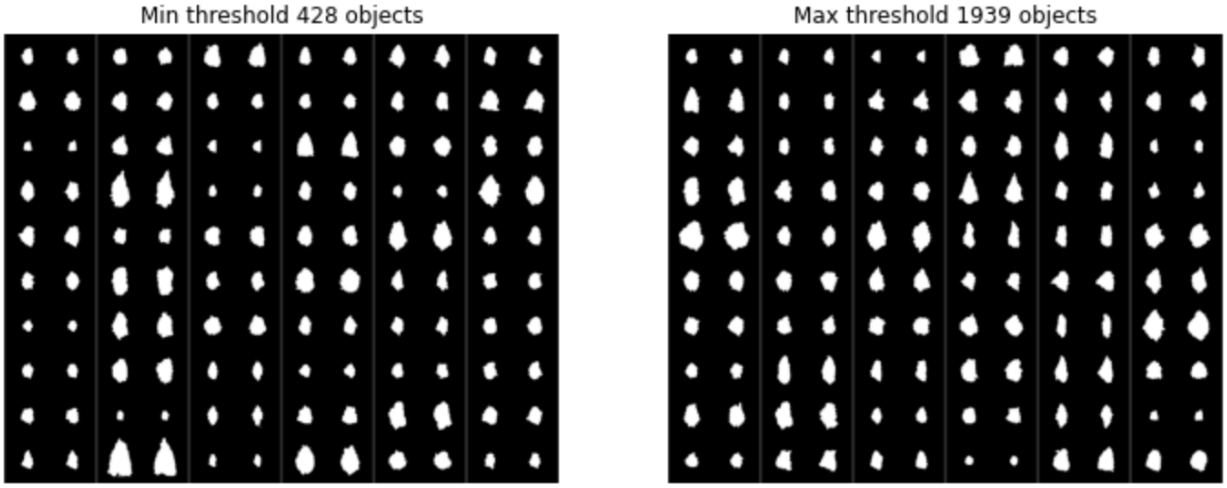

Min threshold 428 objects
Max threshold 1939 objects

As discussed in Methods, this approach for identifying semantically-similar objects is approximate. We aim only to have a low false positive rate (which is what the figures show), but we cannot guarantee a low false negative rate.

#### O2-VAE embedding errors

Why does O2-VAE still have embedding errors? We investigate with the following figure.

**Supplementary Figure 1g: Investigating the cause of O2-VAE embedding errors**
Look at 'high-confidence-image-pairs' where O2-VAE had embedding errors for the threshold $t = 100$ that was chosen for experiments. **a** Sample image pairs. We show either 20 pairs (for two cell lines), or all of the pairs where there were fewer than 20 errors (for three cell lines). **b** Histogram of the 'kNN distance' between the images on a log scale, so that we can tell how far above threshold these image pairs are. **c** The same histogram as b of the 'kNN distance', but for the prealign-VAE.

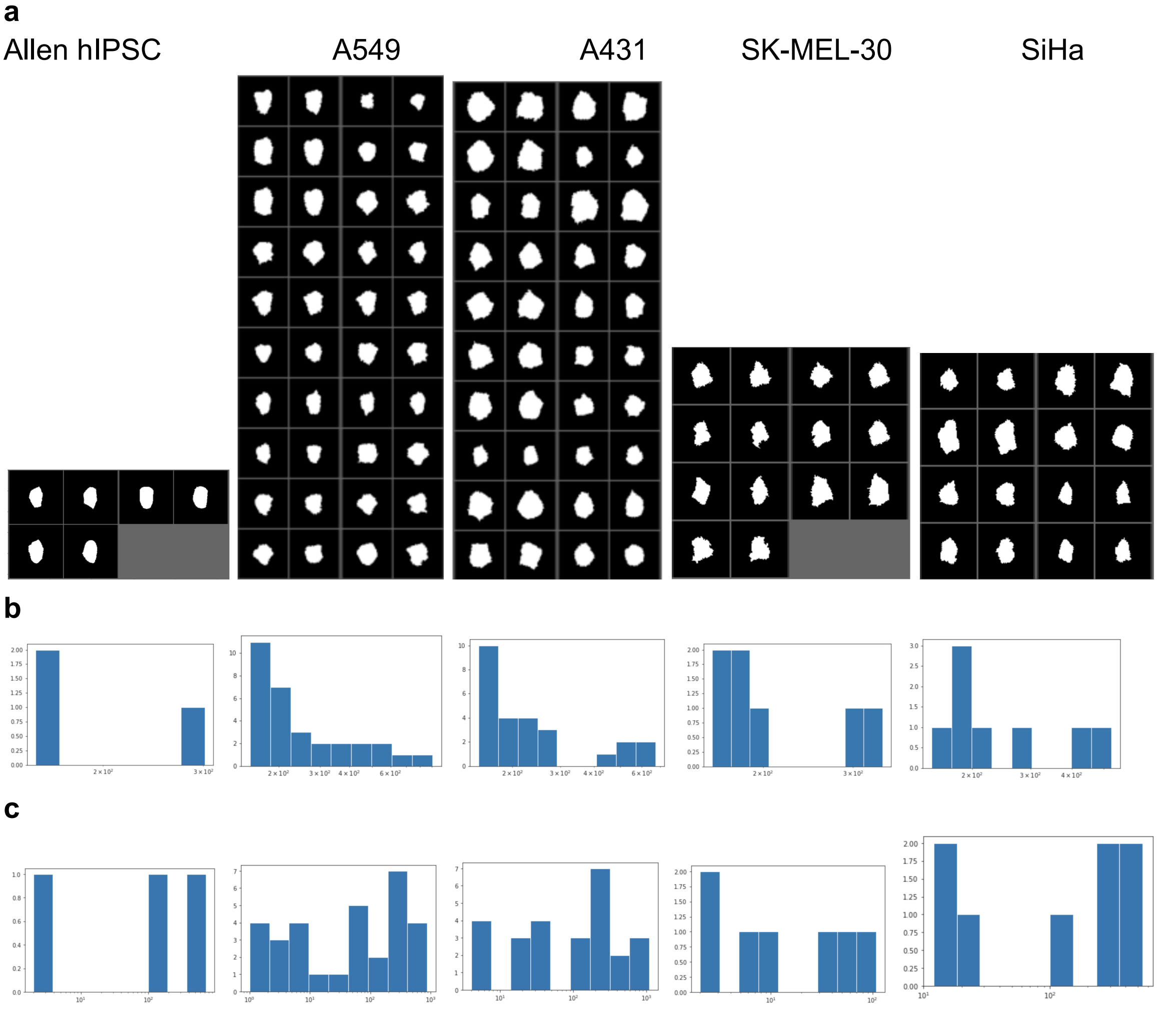


Discussion of Supplementary Figure 1g:

- Firstly note that for 3 of the datasets, there are very few examples: fewer than the 20, so O2-VAE makes very few embedding errors.
- One explanation is that the images may not actually be that visually similar. The definition of “similar” is pixel-based and based on a threshold, so you would expect a few images that are not actually semantically similar. In that case, it’s reasonable for the embedding distance to be larger.
    - One piece of evidence supporting this is to also look at the `k` distribution histograms for prealign-VAE. For some datasets, the distribution of `k` for prealign-VAE is wider. (The cases where this was not true were dataset having very few instances of O2 errors.).
    - But there are few such examples, suggesting that the ‘similar image pair’ metric does achieve the desideratum of having low false positives rate.
- The second explanation is that the images are still close, but the kNN distance is slightly above the threshold. This is also supported by looking at the histogram of kNN values.

### Supplementary 2

This supplementary supports the third Results section.

#### Synthetic shape dataset: PCA of embedding space classes

**Supplementary Figure 2a: PCA of synthetic shape dataset colored by generative factors**

**a** Projection to the plane with PC1 (x-axis) and PC2 (y-axis) coloured by eccentricity (left) and randomness (right). **b** the same, except projection on the plane of PC3 and PC4.

**a**

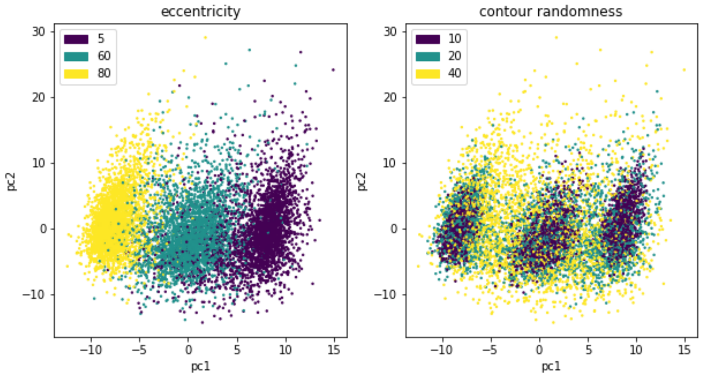


**b**

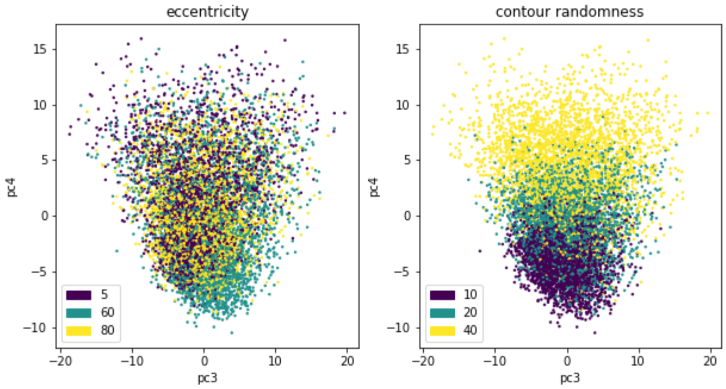


In Fig.3.a (from the main text) we showed samples from the synthetic shape dataset, a distance matrix between class centroids and UMAPs. We used these to claim that eccentricity variation was the dominant factor of variation followed by contour randomness. Here, supplementary Figure 2a shows the same thing. Eccentricity dominates in PC1, and randomness dominates in PC3.

#### Synthetic shape dataset: class overlap for high randomness classes

**Supplementary Figure 2b: image samples for high random classes**

Each row is random samples from each of the three ‘high random’ data classes. The classes are ordered from highest to lowest eccentricity.

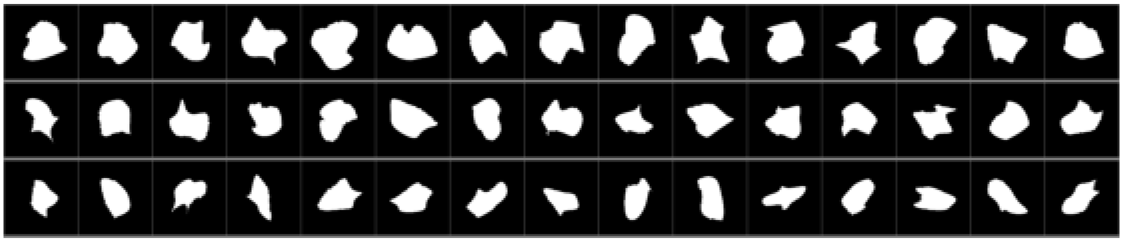

In the third Results section, we claimed that for high randomness classes, the ‘boundary’ between classes with different eccentricity can be overlapping. This can be seen by observing that some samples between classes are very similar, for example the first two rows of the second column. This is a result of the sampled generative process.

The embedding space dimensionality reduction plots also indicate that the high randomness classes are overlapping: UMAP in Fig 3a and PCAs in Supplementary 2a. Since Supplementary Fig.2b shows that the classes really do overlap, we can see that this is likely a feature of the data, and not an issue with the learned representations failing to separate classes.

#### Verification tests: comprehensive and quantitative orientation invariance test

We do a more direct and comprehensive test for orientation invariance, and in Supplementary Figure 2c below, we visualise what these metrics mean

- For each object in the test set, take a set of rotations and flips (Supplementary Figure 2c, left), and compute their representations.
- The representations should not change, but due to discretisation artefacts, the output encodings will change a bit (see Methods discussion). We want to validate that the difference is small. Supplementary Figure 2c (top-right) is a histogram of the distances between encodings of different orientations. We measure the *biggest* embedding distance of the same image due to rotations and flips (the biggest value in Supplementary Figure 2c, top right).
- We measure the embedding distance between the original image and *every other image* in the dataset (the histogram of these values is Supplementary Figure 2c, bottom right). We identify the *smallest* such distance, which is the 1-nearest neighbour (we mark this as a red line in Supplementary Figure 2c, bottom-right).
- We check that the *biggest* encoding distance due to rotations and flips is smaller than the *smallest* encoding distance between other images in the dataset. The top and bottom of Supplementary Fig 2.c have the same x-axis, which shows that this condition is met.

**Supplementary Figure 2c: visualisation of the metrics for quantitative orientation invariance test**

(Left) ten orientations of the same image for testing: five rotation sets for two flips. We take the embeddings for all these images. (Right, top) histogram of the distance matrix values between all ten embeddings. (Right bottom) histogram of the distances between the original image embeddings and all other values, with a red line marking the smallest value. If we have orientation invariance, this red line should be at a larger value than all of the points in the top histogram.

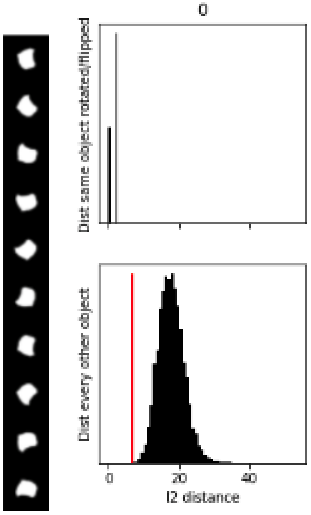


Our invariance test passes for every object in the Allen hiPSC dataset.

#### hiPSC data: extended analysis figures Allen hiPSC cells and nuclei

In the third results section and Fig.3b, we visualise the shape variation in the Allen hiPSC dataset after scale-normalisation. We did clustering, and now show another summarisation approach: dimensionality reduction

**Supplementary Figure 2d: Allen hiPSC scaled data - PCA traversals**
**a** PCA traversals of scale-normalised Allen hiPSC cells. Each image is a point in the representation space visualised by passing that representation through the decoder. Row i is the i'th PC. The mid-point is the origin, and the images in the rest of the row are equally spaced points between -2σ and 2σ where σ is the standard deviation of the representations along that PC. **b** The same PCA traversal plot for the nucleus.

**a**

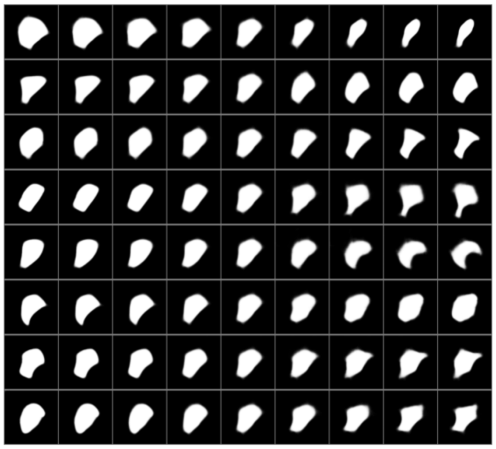

**b**

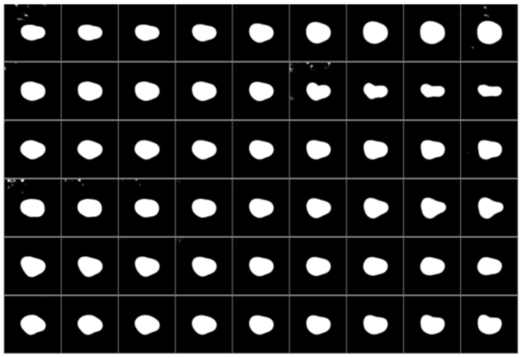

**Supplementary Figure 2e: Allen hiPSC scaled data - UMAP representation**

**a** Scatterplot of UMAP-reduced Allen cell hiPSC cell data (scaled). The colouring is with respect to area. **b** The same UMAP data but we sample images from the real dataset. We create a grid in the UMAP space and sample the image whose reduced embedding is nearest the centroid.

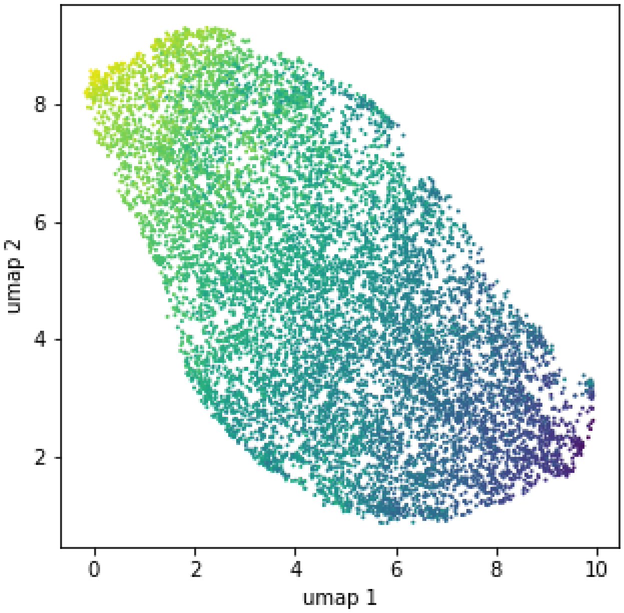


**b**

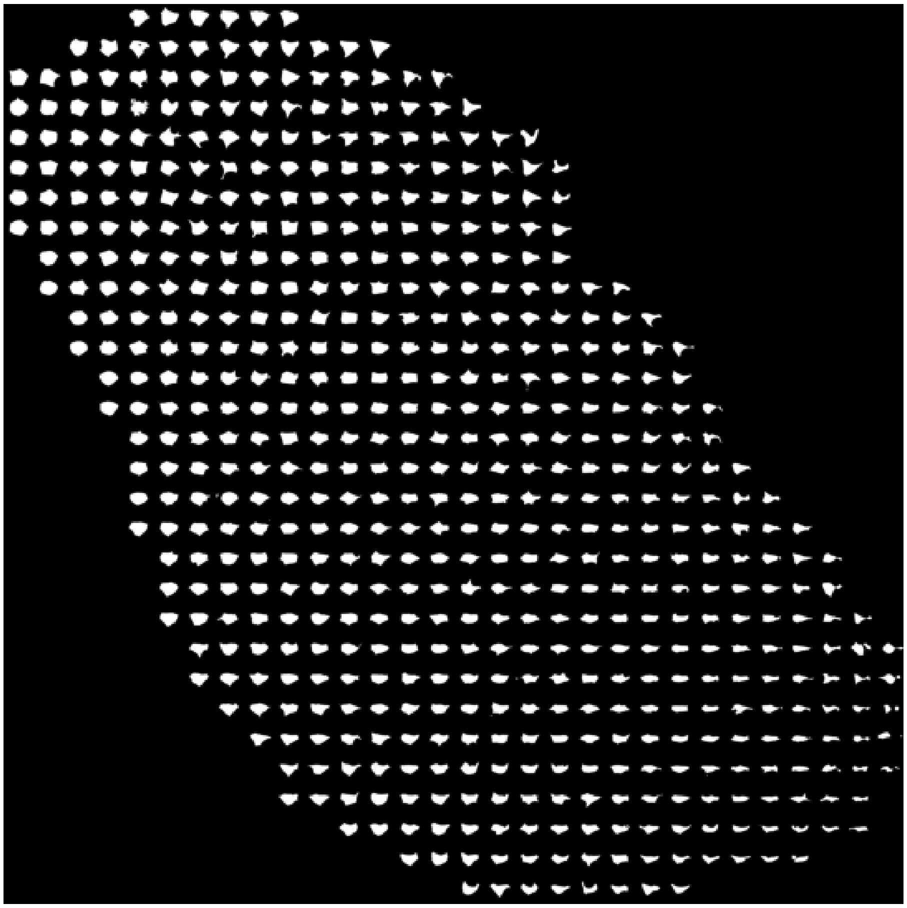

#### hiPSC extended data: visualising shape variation for non-scaled cellular data

In the third results section and Fig.3b, we visualise the shape variation in the Allen hiPSC dataset after scale-normalisation. Here we do clustering and dimensionality reduction of cells without scale normalisation

**Supplementary Figure 2.f: clustering of Allen hiPSCs without scale normalisation**
**a** Cell clustering with GMM and k=14. Prototypes (left), cluster samples (middle) and population frequencies (right).

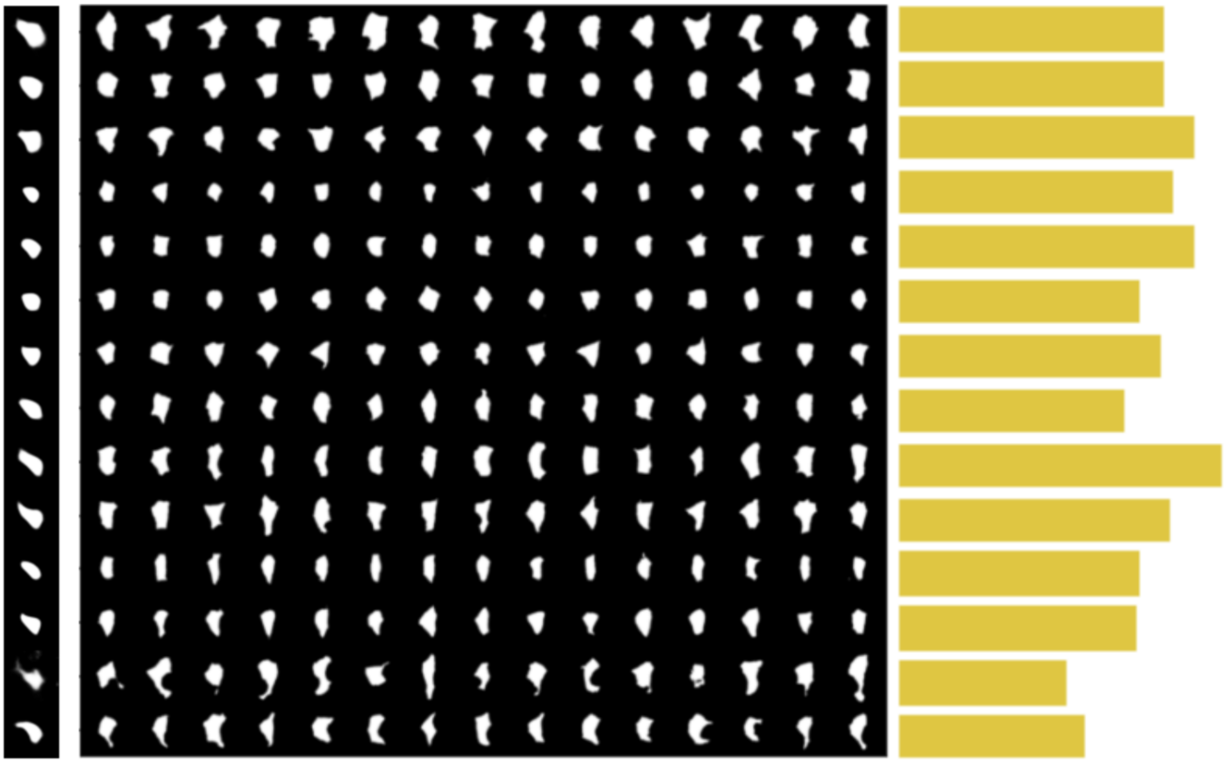

**Supplementary Figure 2g: Allen hiPSC non-data - PCA traversals**
**a** PCA traversals, as described in Supplementary Figure 2d.

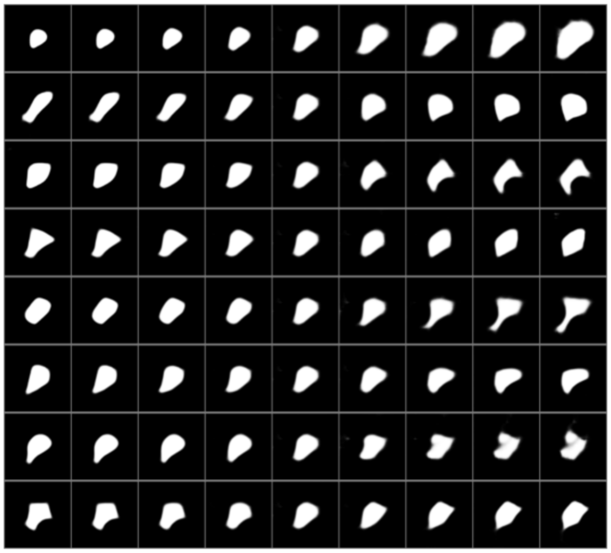

**Supplementary Figure 2h: Allen hiPSC non-scaled cells - UMAP representation**
UMAP-reduced data we sample images from the real dataset

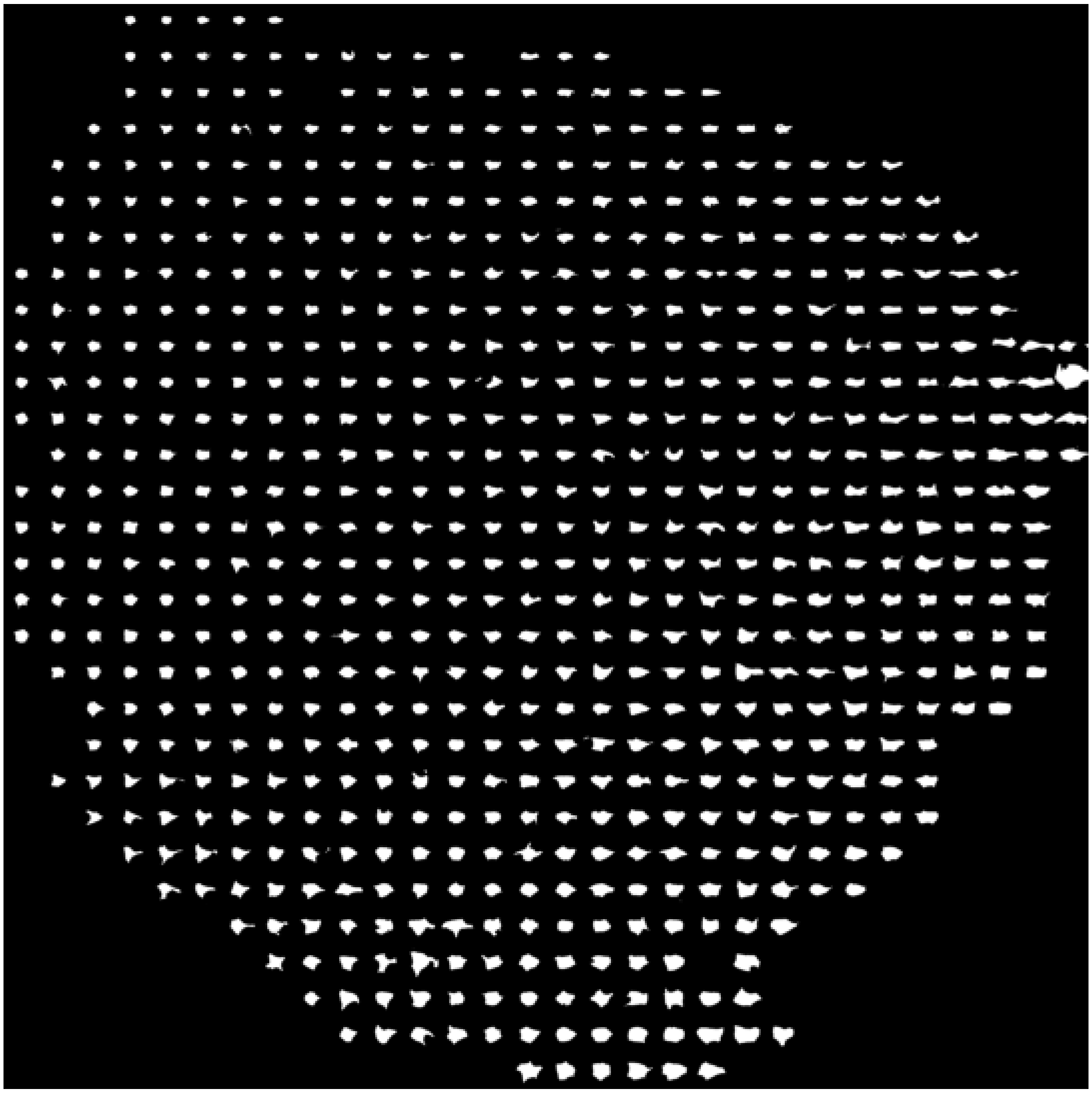

### Supplementary 3

This supplementary supports the fourth Results section.

#### MEFs nucleus clustering

We show the clustering results for nuclei in MEFs to support the claim that LMNA deficient cells have lower prevalence of circular shape groups

**Supplementary Figure 3a: clustering of MEFs nuclei**
Nucleus clustering with GMM and k=8. Prototypes (left), cluster samples (middle) and population frequencies (right).

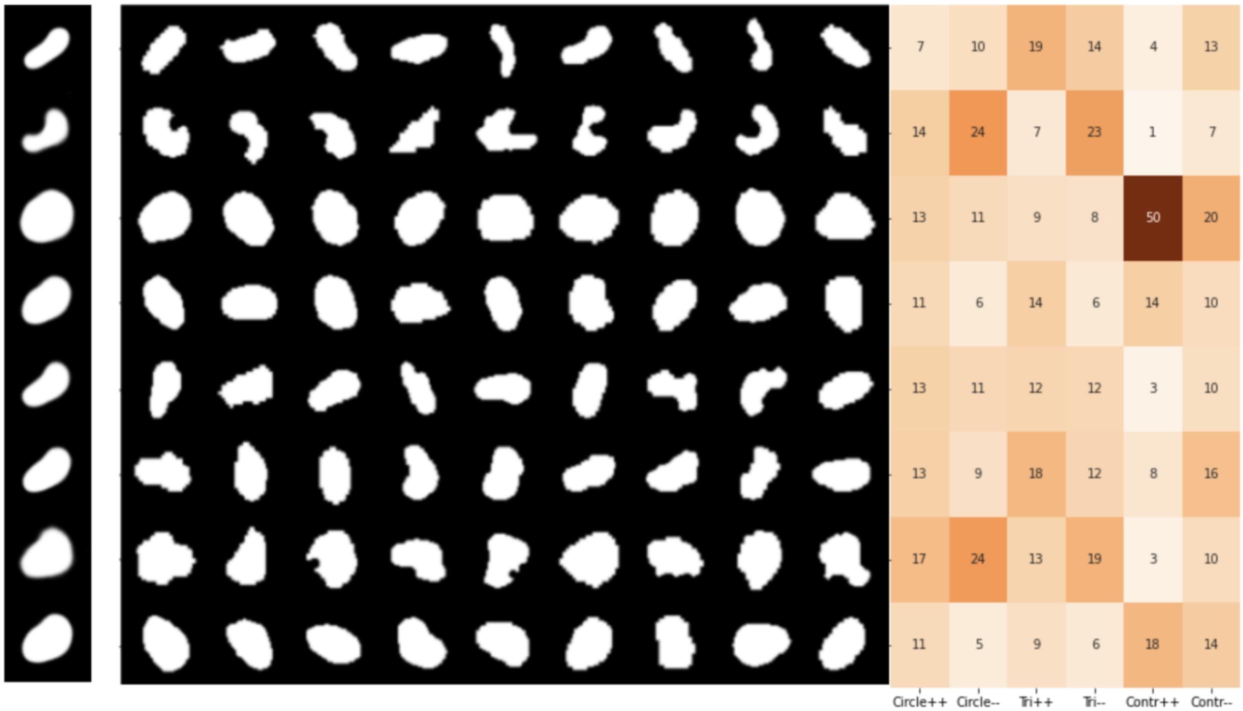


#### Unsupervised mitosis detection in hiPSCs

Using the simple approach described in Results we show further details of the unsupervised approach for mitosis detection.

**Supplementary Table 3a: class-level scores for unsupervised mitosis detection**
For each mitosis state, accuracy scores for whether the cells in that state were detected as 'in mitosis' under the unsupervised detection model described in Results.

| Stage | # samples | Accuracy |
|---|---|---|
| Interphase | 9672 | 99.37% |
| Prophase | 99 | 42.42% |

| | | |
|---|---|---|
| Early prometaphase | 82 | 100.00% |
| Prometaphase / metaphase | 216 | 99.54% |
| Anaphase / telophase paired | 63 | 100.00% |
| Anaphase / telophase unpaired | 135 | 94.81% |

In Supplementary Table 3b, half of the prophase states are missed, but all other states are discovered with more than 90% accuracy. We go deeper into these results with examples in the next figure

**Supplementary Figure 3b: prophase classes detected correctly and incorrectly**
**(**left) Sample prophase cells that were correctly classified as "in mitosis" by our approach. (Right) prophase cells incorrectly classified as "normal" by our model. (the ground truth labels were generated by 3d regular images).

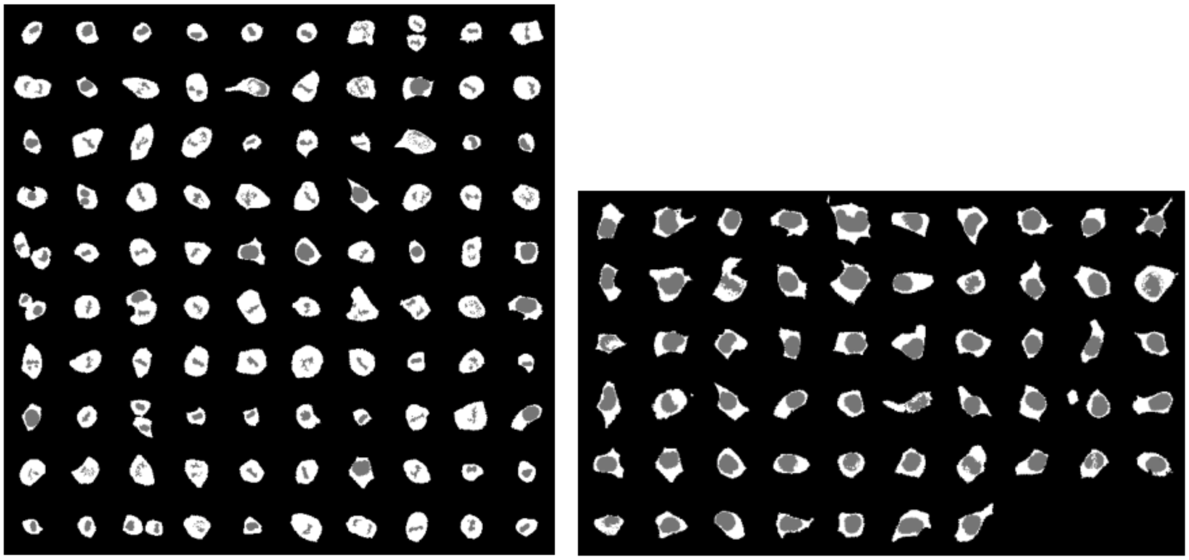

Note, in Supplementary Fig.3b, that most of the prophase cells incorrectly labelled as normal by our approach really do look normal - many look like interphase cells. Their prophase classification is likely based on either more details from 3D images, more details from grayscale information, or the given label could be an error.

Finally, our method labels some cells as outliers (mitosis) that are labelled as 'normal' in the Allen cell collection. We show what these cells look like in Supplementary Fig.3c, and suggest that some of them *may* be mitosis outliers.

**Supplementary Figure 3c: candidate 'missed' mitosis cells**
Sample cells that our unsupervised approach classified as 'mitosis' that were classified as normal in the dataset labels. If some of these really are in mitosis, then these samples illustrate the potential for an unsupervised approach to be a secondary check on other methods.

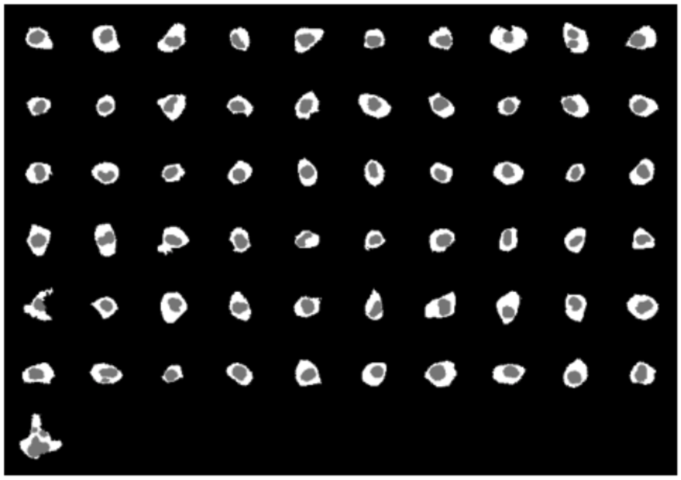

#### Organelle contact rates

In the fourth results section and Fig.4e-f we report on contact rates for mitochondria subgroups. Here is some additional supporting data.

**Supplementary Figure 3d: cluster samples and prevalence for mitochondria groups used in contact rate analysis**

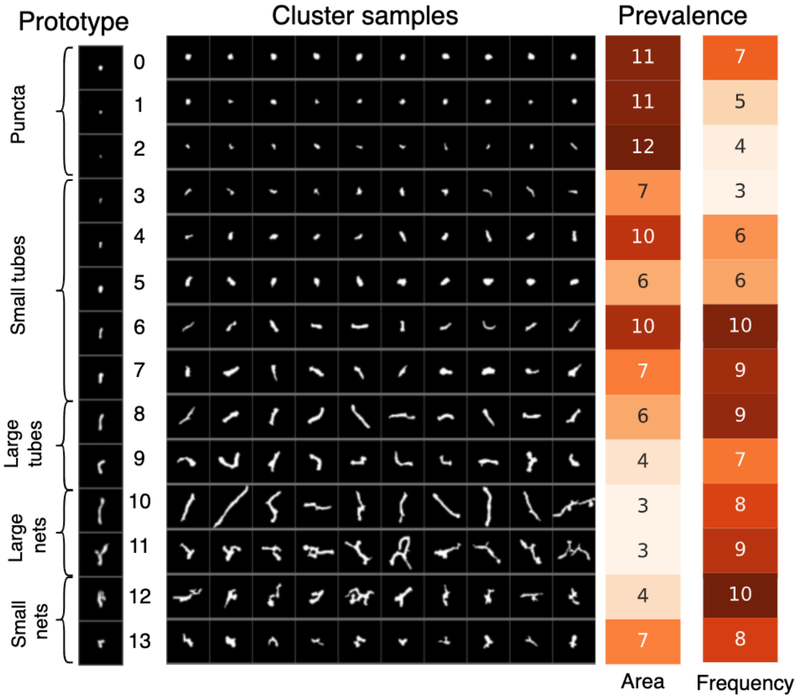


**Supplementary Figure 3e: contact rates from Fig.4g for sub-experiments:**
The sub-experiments from Fig.4g, broken down into 3 subgroups collected as different experiments: ‘exp1’, ‘exp2’, and ‘exp3’. The last heatmap, ‘bodipy’ , is the tag for lipid droplet.

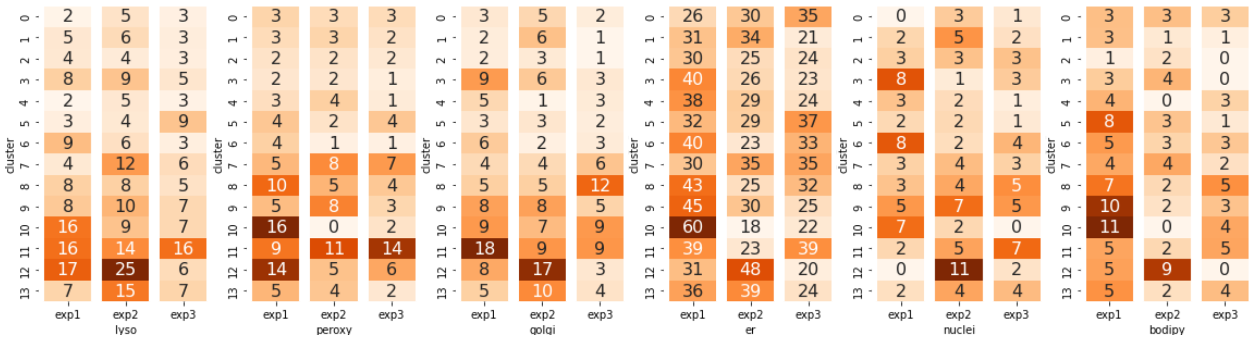
cluster
0 1 2 3 4 5 6 7 8 9 10 11 12 13
2 5 3
5 6 3
4 4 3
8 9 5
2 5 3
3 4 9
9 6 3
4 12 6
8 8 5
8 10 7
16 9 7
16 14 16
17 25 6
7 15 7
exp1 exp2 exp3
lyso
3 3 3
3 3 2
2 2 2
2 2 1
3 4 1
4 2 4
4 1 1
5 8 7
10 5 4
5 8 3
16 0 2
9 11 14
14 5 6
5 4 2
exp1 exp2 exp3
peroxy
3 5 2
2 6 1
2 3 1
9 6 3
5 1 3
3 3 2
6 2 3
4 4 6
5 5 12
8 8 5
9 7 9
18 9 9
8 17 3
5 10 4
exp1 exp2 exp3
golgi
26 30 35
31 34 21
30 25 24
40 26 23
38 29 24
32 29 37
40 23 33
30 35 35
43 25 32
45 30 25
60 18 22
39 23 39
31 48 20
36 39 24
exp1 exp2 exp3
er
0 3 1
2 5 2
3 3 3
8 1 3
3 2 1
2 2 1
8 2 4
3 4 3
3 4 5
5 7 5
7 2 0
2 5 7
0 11 2
2 4 4
exp1 exp2 exp3
nuclei
3 3 3
3 1 1
1 2 0
3 4 0
4 0 3
8 3 1
5 3 3
4 4 2
7 2 5
10 2 3
11 0 4
5 2 5
5 9 0
5 2 4
exp1 exp2 exp3
bodipy

### Supplementary 4

This supplementary supports the fifth Results section.

#### Texture only experiments

**Supplementary Figure 4a: dimensionality reduction of texture only experiment**
**a** 2D UMAP reduction of data trained on a synthetic cell shape dataset with consistent shape, and varying texture, coloured by texture level. The texture classes are separated. **b** PCA reduction (PC1 vs PC2) also coloured by texture. The classes are not well-separated using this linear method.

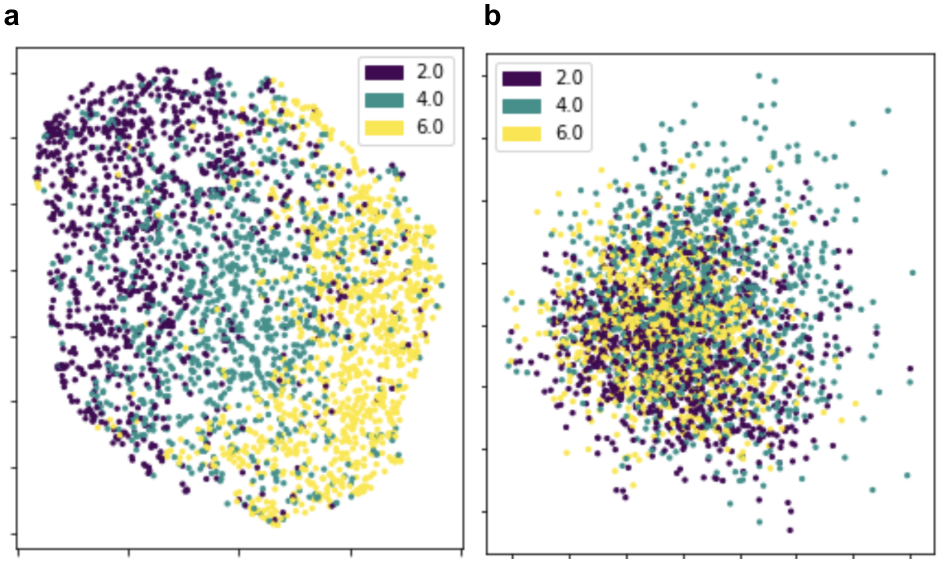


Interestingly, though the UMAP organises the embedding space by texture, the PC1-PC2 plot does not, so the texture variation in embedding space is not linear (we also checked that plotting lower PCs does not reveal the structure). But we do know that the classes really are separated based on the linear probing scores (0.93).

#### Joint texture and shape experiments

Here we vary eccentricity, randomness, and texture (3 levels for each variable for 27 total classes). This extends Fig 5 where we did not model cell contour randomness. Generally we find that separating the classes with this many factors of variation becomes difficult (especially for texture).

**Supplementary Figure 4a: summary representation space visualisations for joint shape and texture experiment**
For the synthetic cell shape dataset, UMAP reductions are colored by **a** eccentricity, **b** contour randomness, and **c** texture. **d** distance matrix between the class centroids. There is a 3-level hierarchy of classes. The lowest is texture, so the first 3 classes are (low, medium, high) texture.

The mid-level is eccentricity, so the first 3 cells are low contour randomness, then the next 3 are medium contour randomness and so on. The highest level is eccentricity.

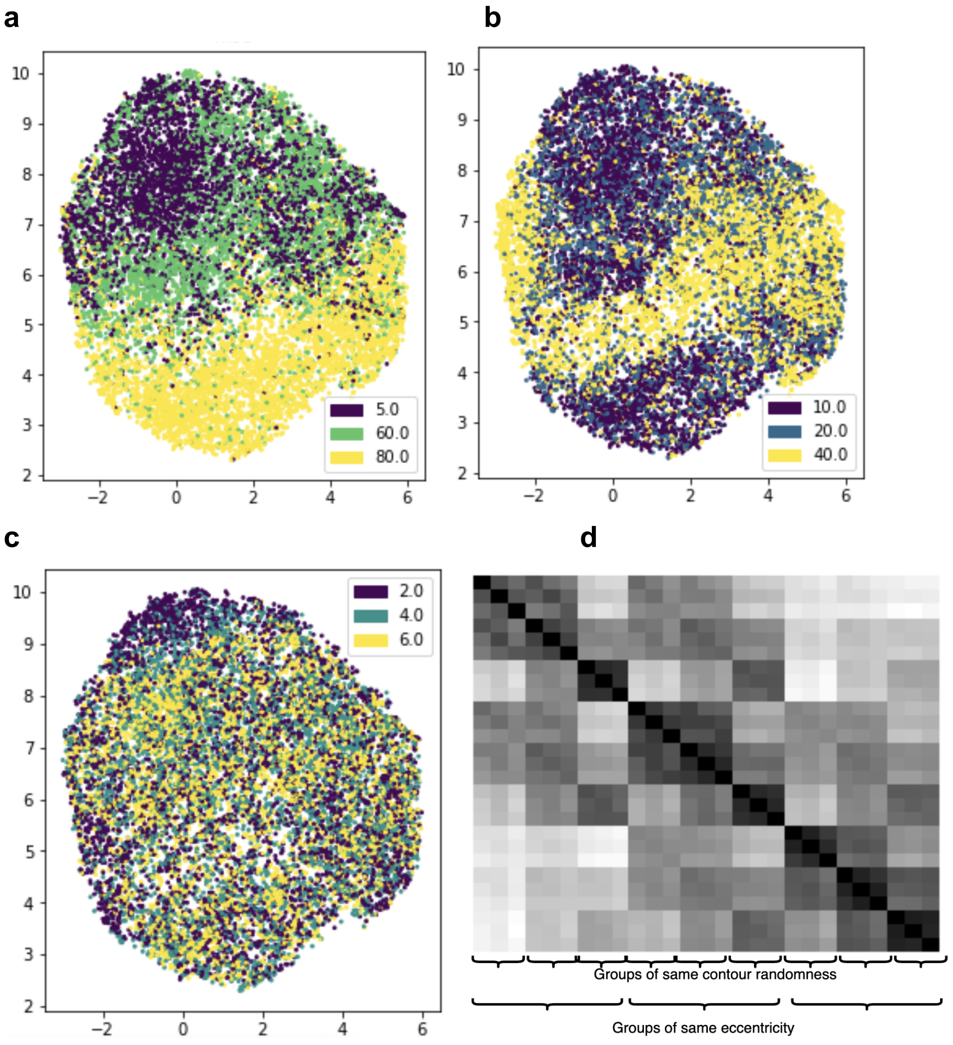


First, these results show that there is a hierarchy for which factors of variation vary the most in representation space: eccentricity is the most dominant, then contour randomness, then texture. Second, texture features are not as well-separated in embedding space compared with the shape features, suggesting room for improving the model.
